# TreeSwift: a massively scalable Python tree package

**DOI:** 10.1101/325522

**Authors:** N. Moshiri

**Affiliations:** Department of Computer Science and Engineering, UC San Diego, 92093, USA

**Keywords:** phylogenetics, tree traversal, scalable, Python

## Abstract

Phylogenetic trees are essential to evolutionary biology, and numerous methods exist that attempt to extract phylogenetic information applicable to a wide range of disciplines, such as epidemiology and metagenomics. Currently, the three main Python packages for trees are Bio.Phylo, DendroPy, and the ETE Toolkit, but as dataset sizes grow, parsing and manipulating ultra-large trees becomes impractical for these tools. To address this issue, we present TreeSwift, a user-friendly and massively scalable Python package for traversing and manipulating trees that is ideal for algorithms performed on ultra-large trees.

## 1. Motivation and significance

Phylogenetic trees are essential to evolutionary biology, and phylogenetic methods are applicable to a wide range of disciplines, such as epidemiology [1, 2] and metagenomics [3, 4, 5]. However, the datasets analyzed by these methods are growing rapidly as sequencing costs continue to fall, emphasizing the need for scalable methods of tree traversal and manipulation. Beyond the analysis of real datasets, phylogenetic approaches can be utilized in the analysis of potentially massive datasets generated by simulation experiments [6].

Methods for performing phylogenetic analyses such as clustering [7] and rerooting [8] are typically presented as a series of higher-level tree traversals and manipulations. The developers of these tools do not commonly implement basic tree processing from scratch: they typically utilize existing tree packages to handle low-level tasks and instead implement their algorithms as a series of calls to functions of these packages. As a result, the performance of such a tool depends not only on the time complexity of its algorithm, but also on the performance of the underlying tree package.

Currently, the three main Python packages for trees are the Bio.Phylo module of Biopython [9], DendroPy [10], and the ETE Toolkit [11]. The three tools are simple to integrate into new methods, include a plethora of functions that cater to most phylogenetics needs, and are fast for reasonablysized trees. However, as dataset sizes grow, parsing and manipulating ultralarge trees becomes impractical. We introduce TreeSwift, a scalable cross-platform Python package for traversing and manipulating trees that does not require any external dependencies, and we compare its performance against that of Bio.Phylo, DendroPy, and the ETE Toolkit.

## 2. Software description

### 2.1. Software overview

TreeSwift is a pure-Python package that has no required external dependencies and which has been tested on Python versions 2.6–2.7 and 3.3–3.7. It is also compiled and hosted on PyPI, meaning it can easily be installed with a single pip command without any need for administrative privileges or any advanced knowledge. This is essential to contrast against the current state-of-the-art, ETE Toolkit, which requires the Six and NumPy Python libraries to install if the user has administrative privileges or Anaconda/Miniconda to install if the user doesn’t, and BioPython, which requires a C compiler and the NumPy Python library as well as the computer fluency to compile tools from source using a `Makefile`.

A key feature of TreeSwift is its simplicity in class design in order to reduce time and memory overhead of loading, traversing, and manipulating trees. The entire package consists of just two classes: a `Node` class, which contains the data and local relationships, and a `Tree` class, which handles manipulation and traversal on the `Node` objects. A key distinction between TreeSwift and DendroPy is that DendroPy stores bipartition information to enable efficient comparisons between multiple trees that share the same set of taxa, but because TreeSwift is designed for the fast traversal and manipulation of individual trees (and not for the comparison of multiple trees), TreeSwift forgoes this feature to avoid the accompanied overhead, resulting in a much lower memory footprint and faster execution of equivalent functions (Fig. 1).

**Figure 1:**
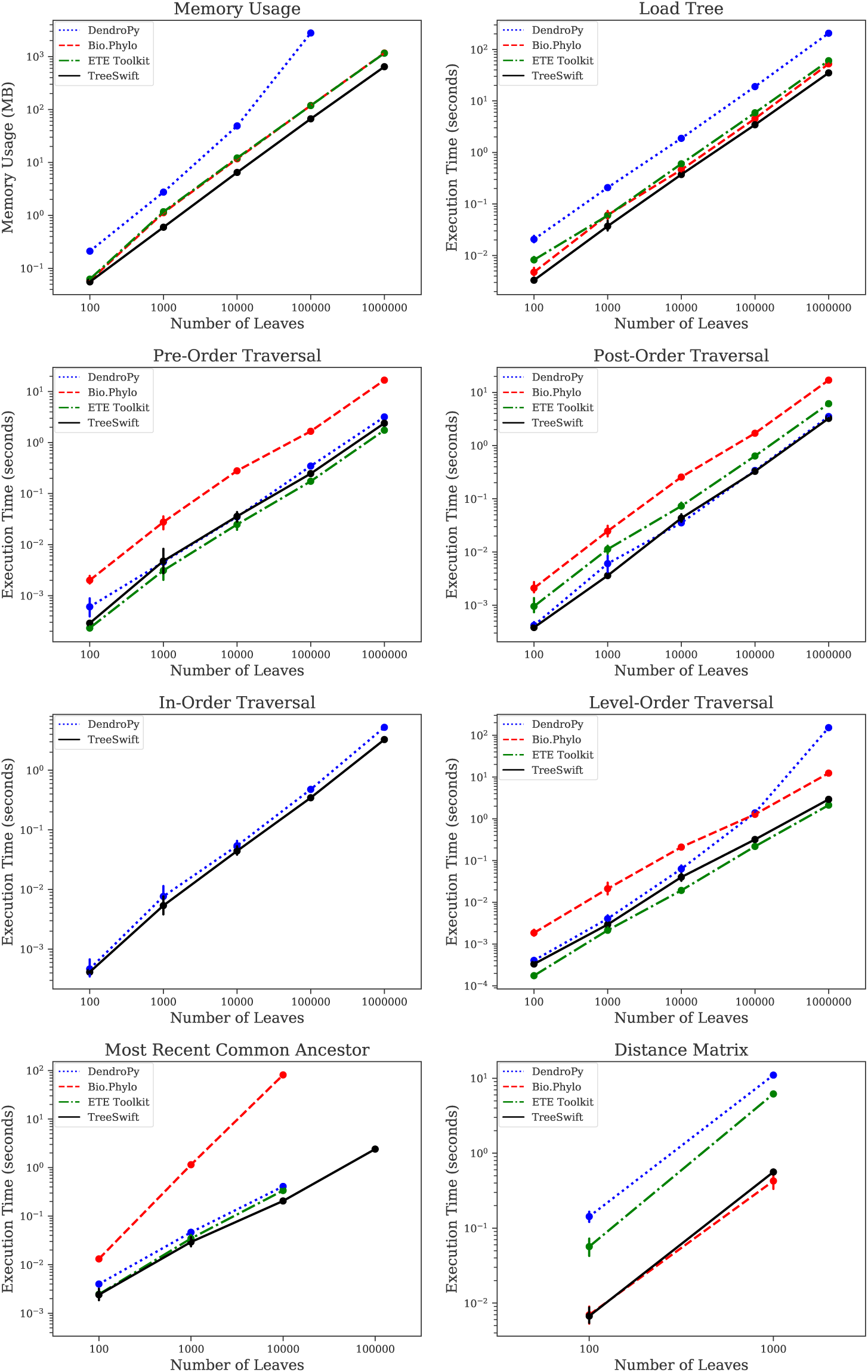
Runtimes of DendroPy, Bio.Phylo, the ETE Toolkit, and TreeSwift for a wide range of typical tree operations using trees of various sizes, as well as memory consumption after loading a tree (see Sec. 3 for details).

### 2.2. Software functionalities

TreeSwift supports loading trees in the Newick, Nexus, and NeXML file formats via the `read tree newick, read tree nexus`, and `read tree nexml` functions, respectively. Inputs to these functions can be strings, plaintext files, or gzipped files, and TreeSwift handles the nuances of parsing them internally to maintain user-friendly operability.

TreeSwift provides generators that iterate over the nodes of a given tree in a variety of traversals, including pre-order, in-order, post-order, level-order, and root-distance-order. TreeSwift also allows for the modification of the structure of a given tree by simply modifying the `Node` objects of the tree. These built-in generators and modifiers intend to provide developers a simple yet efficient manner in which to implement their own algorithms such that they only need to consider higher-level details of the traversal process.

TreeSwift also provides the ability to compute various summarizing statistics of a given tree, such as tree height, average branch length, patristic distances between nodes in the tree, treeness [12], and the Gamma statistic [13]. Beyond numerical statistics to describe trees, TreeSwift can also generate a visual summary of a tree in the form of a Lineages-Through-Time (LTT) plot [14], a feature not currently implemented in any other Python tree package.

## 3. Benchmarking

TreeSwift, Bio.Phylo, DendroPy, and the ETE Toolkit were benchmarked by running the following operations on binary trees with 100, 1,000, 10,000, 100,000 and 1,000,000 leaves: pre-order traversal, post-order traversal, in-order traversal, level-order traversal, finding the most recent common ancestor (MRCA) of all pairs of leaves, and computing the pairwise distances of all pairs of leaves. These operations were chosen because they are fundamental algorithms commonly performed on tree structures.

Timing was performed on a computer running CentOS release 6.6 (Final) with an Intel(R) Xeon(R) CPU E5-2670 0 at 2.60GHz and 32 GB of RAM. Each point shows the average and 95% confidence interval over 10 runs. All scripts and data can be found in the following GitHub repository: github.com/niemasd/TreeSwift-Paper

As can be seen in Figure 1, for all tests, TreeSwift consistently outper-forms the existing tools. When loading trees, TreeSwift is consistently faster than the other tools, and it consumes significantly less memory; DendroPy is consistently around an order of magnitude slower and consumes over an order of magnitude more memory than TreeSwift. When performing pre-order traversals, the ETE Toolkit, DendroPy, and TreeSwift have similar runtimes, with the ETE Toolkit seeming to run slightly faster; Bio.Phylo is close to an order of magnitude slower. When performing post-order traversals, DendroPy and TreeSwift have similar runtimes; the ETE Toolkit is consistently around half an order of magnitude slower, and Bio.Phylo is consistently an order of magnitude slower. When performing in-order traversals, DendroPy and TreeSwift have similar runtimes, with TreeSwift seeming to run slightly faster; Bio.Phylo and the ETE Toolkit did not appear to have in-order traversals implemented. When performing level-order traversals, the ETE Toolkit and TreeSwift have similar runtimes, with the ETE Toolkit seeming to run slightly faster; Bio.Phylo is consistently around an order of magnitude slower, and while DendroPy is similar in runtime for trees with up to 10,000 leaves, its runtime significantly worsens when traversing trees with 100,000 and 1,000,000 leaves. When finding the most recent common ancestor, the ETE Toolkit, DendroPy, and Treeswift have similar runtimes for trees with up to 100,000 leaves, with TreeSwift seeming to run slightly faster; Bio.Phylo is consistently over an order of magnitude slower, and TreeSwift was the only tool able to scale this operation to trees with 1,000,000 leaves. When computing a matrix containing the distances between all pairs of leaves, TreeSwift and Bio.Phylo have similar runtimes; the ETE Toolkit and DendroPy are over an order of magnitude slower.

## 4. Illustrative example

In the following example, a tree is loaded from a gzipped file, the minimum distance from each node to a leaf is computed, the minimum leaf distance of the root is printed, and a Lineages-Through-Time (LTT) plot is created (Fig. 2).

**Figure 2:**
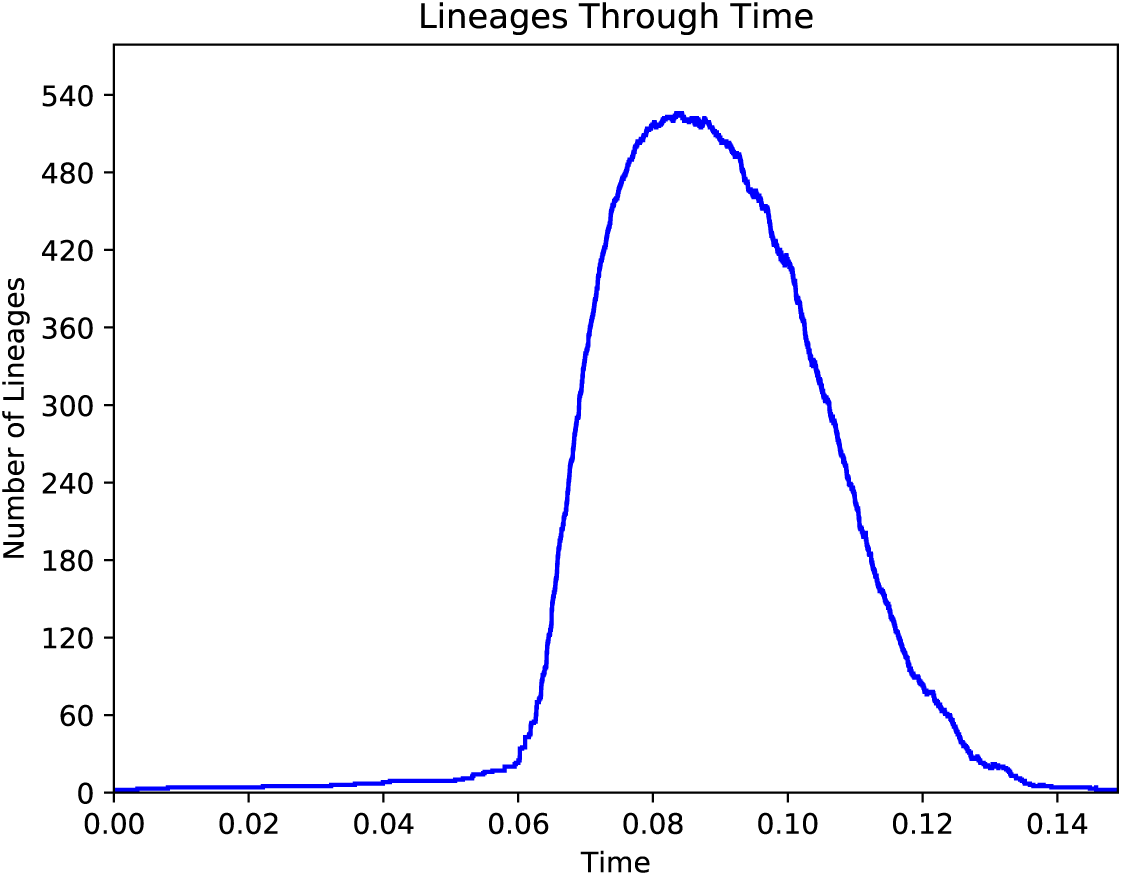
Example Lineage-Through-Time (LTT) plot generated using TreeSwift.

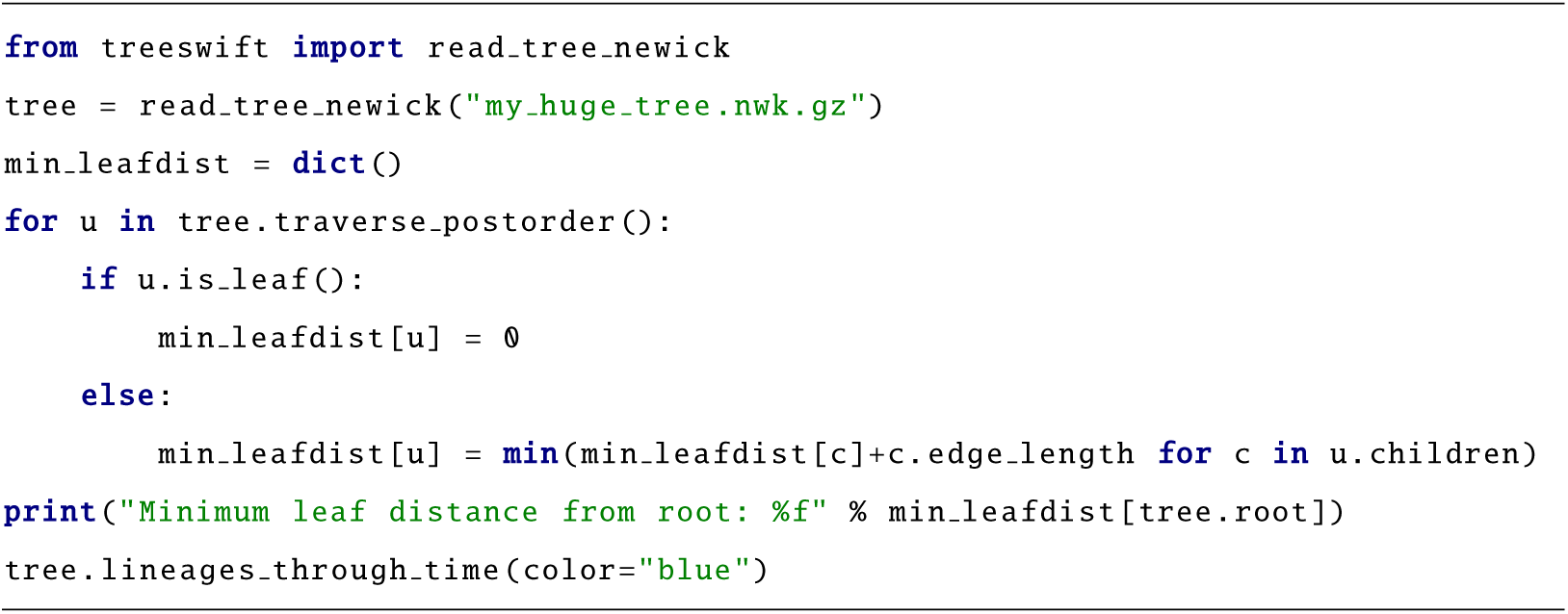

## 5. Impact

The key impact of TreeSwift is its significant performance improvement over existing Python tree packages (Fig. 1). For almost all tested tree operations, TreeSwift performed tasks significantly faster than all existing tools (by orders of magnitude at times), and it was the only tool that not only had all tested functions implemented, but that also was able to scale to the largest of tested datasets. Further, TreeSwift’s memory consumption was significantly lower than all existing tools. Thus, phylogenetic tools written in Python can utilize TreeSwift for scalability.

Further, TreeSwift was designed to be simple to use. As can be seen in the example code in Section 4, a user with minimal Python experience can generate a Lineages-Through-Time (LTT) plot in just 3 lines of Python code. Even complex tree algorithms can be implemented cleanly by utilizing TreeSwift’s traversal generators [7].

It must be emphasized that, although TreeSwift was designed with the field of phylogenetics in mind, the package is general in that it can be utilized with any arbitrary tree structure, including those in non-phylogenetic applications [15]. Thus, its utility can extend well beyond its intended phylogenetics audience.

## 6. Conclusions

In this article, we presented TreeSwift, a pure-Python package for loading, traversing, and manipulating trees in a massively-scalable manner. The current version implements a wide range of typical tree operations, and due to its simple design, it has significant room for developers from other disciplines to further expand its capabilities to target a larger suite of potential applications.

## Acknowledgements

This work was supported by NIH subaward 5P30AI027767-28 to NM. We would like to acknowledge Siavash Mirarab for his mentorship. We would also like to acknowledge Jeet Sukumaran and Mark Holder, as DendroPy provided much motivation during TreeSwift’s development.

## Required Metadata

### Current code version

Ancillary data table required for subversion of the codebase. Kindly replace examples in right column with the correct information about your current code, and leave the left column as it is.

**Table 1:** Code metadata (mandatory)

| Nr. | Code metadata description | Please fill in this column |
| --- | --- | --- |
| C1 | Current code version | 1.1.3 |
| C2 | Permanent link to code/repository used for this code version | github.com/niemasd/TreeSwift |
| C3 | Legal Code License | GNU GPL v3.0 |
| C4 | Code versioning system used | git |
| C5 | Software code languages, tools, and services used | Python |
| C6 | Compilation requirements, operating environments & dependencies | Python, pip (optionally matplotlib for LTT plot visualization) |
| C7 | If available Link to developer documentation/manual | github.com/niemasd/TreeSwift/wiki |
| C8 | Support email for questions |  |

